# Proteomic insights into a *M. tuberculosis* clinical isolate with an increased propensity to form viable but non-replicating subpopulations during acid stress

**DOI:** 10.1101/2025.10.03.680234

**Authors:** Nastassja L. Kriel, Julian Coetzee, Jacoba M. Mouton, Samantha Sampson

## Abstract

Phagosome acidification is one of the challenges faced by *Mycobacterium tuberculosis* during infection. This intracellular pathogen is known to adapt to its stressful environment through stress response pathways and by secreting proteins to modify the host immune response for survival and proliferation. However, *M. tuberculosis* also holds the potential to form viable but non-replicating (VBNR) and antibiotic tolerant persisters in response to environmental stress, including acid stress. In this study we used a *in vitro* acid stress model to stimulate the formation of a VBNR subpopulation in a *M. tuberculosis* clinical isolate with an increased propensity to form VBNR bacteria. Mass spectrometry-based proteomics was used to characterize the cellular proteome and culture filtrate proteome of actively replicating (pH 6,5) and VBNR enriched (pH 4,5) cultures. We show that in response to acid stress, *M. tuberculosis* S169 increases the expression of known stress response proteins, including the methyltransferase Rv1405c and the acid stress response two-component regulatory protein TcrX. Interestingly, we found that the dormancy response regulon components were less abundant in acid stressed *M. tuberculosis* S169. Our protein aggregation capture culture filtrate proteomic approach revealed that the culture filtrates of low pH stressed *M. tuberculosis* S169 contained less proteins than that of actively replicating cultures. We identified several proteins previously implicated in *M. tuberculosis* persistence, including toxin-antitoxin proteins (VapC51 and VapB10), the chorismate mutase (Rv1885c), and several uncharacterized proteins. The observed differences identified in the characterisation of this clinical isolate in comparison to published *M. tuberculosis* H37Rv highlights the need to investigate *M. tuberculosis* clinical isolates for a more representative understanding of the tuberculosis stress response.

**Author Summary:** Tuberculosis is caused by *Mycobacterium tuberculosis* and this pathogen can form a subpopulation of viable but non-replicating (VBNR) cells that are recalcitrant to antibiotic treatment. These persister bacteria increases the risk of treatment failure and tuberculosis recurrence following treatment. Stimulation of a persister population through triggered persister formation can be achieved by environmental stress factors such as low pH, nutrient starvation, hypoxia, and antibiotic exposure. In this study we investigate the cellular and culture filtrate proteomes of a high persister forming clinical isolate, *M. tuberculosis* S169, in response to acid stress. We show that following the stimulation of a VBNR subpopulation in response to acid stress, several known acid stress response proteins are more abundant in VBNR enriched cultures. Interestingly, we found that stress response proteins were less abundant. Using a protein aggregation capture approach we successfully characterized culture filtrates *M. tuberculosis* cultures, reducing the bacterial culture amount required for these experiments. Culture filtrates differed between actively replicating and VBNR enriched cultures. Several immunogenic proteins were identified in a higher abundance in the culture filtrates of VBNR enriched cultures.

## Introduction

*Mycobacterium tuberculosis*, the pathogen which causes tuberculosis (TB), primarily infects macrophages. During infection, *M. tuberculosis* is exposed to high levels of reactive oxygen and nitrogen intermediates, reduced oxygen and nutrient availability, and low pH (1). *M. tuberculosis* can replicate under these conditions or persist in a viable, but non-replicating (VBNR) state (2). Persister *M. tuberculosis*, can coexist with replicating mycobacteria during infection, but these bacteria are transiently insensitive to antibiotic treatment due to their non- or slow-replicating nature (2,3). These antibiotic recalcitrant bacteria contribute to the length of TB treatment regimens, where multiple antibiotics are required to treat infections for extended periods of time (4,5). Persister subpopulations may resume growth under favourable conditions, increasing the possibility for recurrent disease following treatment (6,7).

Bacterial persisters are thought to arise from spontaneous persistence or triggered persistence (8). Spontaneous persistence describes the stochastic formation of persisters at a rate that is constant during growth, accounting for approximately 1% of the bacterial population in stationary phase (9,10). Triggered persistence refers to the formation of a persister subpopulation in response to a stress signal such as starvation, population density, pH stress, immune factors and drug treatment (8). In other bacterial species, mechanisms of persister formation include metabolic slow down, modulation of nucleoid-associated proteins, expression of drug efflux pumps, activation of toxin-antitoxin modules, and upregulation of stress response genes (11–15). Similarly, a multiple stress dormancy model for *M. tuberculosis* induced lower energy metabolism, reduced transcription and translation, and increased expression of stress response genes as mechanisms of dormancy (2). Several studies have highlighted the upregulation of the dormancy response regulon (DosR) in response to low oxygen and nitric oxide exposure, further emphasizing the role of stress response proteins in bacterial dormancy (16–18). Toxin-antitoxin modules were also more abundant in the proteomes of nutrient starved *M. tuberculosis*, highlighting the cross-species similarities in persister linked pathways (19).

The intracellular pathogen *M. tuberculosis* interacts with the host by presenting various molecules on its cell surface, but also through the secretion of molecules (20,21). *M. tuberculosis* secretion substrates play a role in nutrient acquisition, host epigenetic modification, prevention of phagosome maturation, modulation of cytokine response, autophagy, redox regulation, and necrosis and bacterial dissemination (20,22). Identification of proteins secreted by *M. tuberculosis*, especially in response to environmental stress, is crucial in understanding how *M. tuberculosis* subverts killing by host macrophages (23,24). *Salmonella* persisters have been shown to not only maintain a metabolically active state, but to secrete effector proteins which reprogrammed the host cell polarization form a pro-inflammatory to anti-inflammatory state (25). Furthermore, advances in immunopeptidomics have highlighted the need to investigate proteins identified in the extracellular region of *M. tuberculosis* by demonstrating that ESX secretion substrates are the prominent source of MHC-I presented *M. tuberculosis* peptides (26). This finding highlights the importance of investigating *M. tuberculosis* secreted proteins as anti-TB vaccine development candidates.

In this study we sought to characterize both the cellular proteome and the culture filtrate of a *M. tuberculosis* clinical isolate which has an increased propensity to form VBNR subpopulations (7). This clinical isolate, *M. tuberculosis* S169, was obtained from an HIV-negative patient who failed treatment following standard 6-month anti-TB treatment (27). *M. tuberculosis* S169 is susceptible to anti-TB treatment and no drug-resistance conferring mutations were identified using whole genome sequencing (7). We hypothesize that VBNR *M. tuberculosis* may secrete a different subset of proteins to that of actively replicating *M. tuberculosis*. Given the abundance of VBNR *M. tuberculosis* in bacterial culture, we opted to use this clinical isolate with an increased propensity to form VBNR *M. tuberculosis*, to increase the likelihood of identifying VBNR secreted proteins (10).

We used a low pH stress model to trigger the formation of a VBNR subpopulation and verified the enrichment of a VBNR subpopulation using a dual replication reporter plasmid (28,29). To characterize the culture filtrates of actively replicating and VBNR enriched cultures, we made use of a protein aggregation capture approach (30). Our culture filtrate mass spectrometry approach required smaller amounts of bacterial culture, reducing experimental time, technical variability from pooling multiple cultures, experimental cost, and in the case of *M. tuberculosis*, biohazardous risk.

## Materials and Methods

### Bacterial culture and acid stress exposure

*M. tuberculosis* clinical isolate S169 obtained from Dr Stephanus Malherbe was collected by the Catalysis TB – Biomarker Consortium, Ethical approval granted by Stellenbosch University Human Research Ethics Committee (registration number N10/01/013) (27). Informed consent was obtained from all study participants (27). All reagents were purchased from Sigma Aldrich unless otherwise stated. All experiments were performed in biological triplicate.

Liquid mycobacterial cultures were grown in Middlebrook 7H9 supplemented with 10% dextrose-catalase (DC), 0.2% glucose and 0.05% Tween-80 (7H9-DC). The clinical isolate was transformed with the replication reporter plasmid pTiGc and transformants were cultured in the presence of 25 µg/ml kanamycin at 37°C. For acid stress, *M. tuberculosis* S169::pTiGc was sub-cultured in 7H9-DC pH 6.5 supplemented with 4mM theophylline to an OD_600_ of ∼1.0 prior to washing with culture media. Cultures were resuspended in fresh 7H9-DC at pH 6.5 and pH 4.5 and incubated at 37°C for 48h. Fluorescence dilution was used for the identification of VBNR mycobacteria as previously described (28). Aliquots of bacterial culture pre- and post-acid stress were plated to verify bacterial viability following acid stress.

### Imaging flow cytometry sample preparation, acquisition, and analyses

Cultured *M. tuberculosis* S169::pTiGc was sonicated for 12 minutes at 36kHz (Zues, Sonicator bath) before filtering through a 40 µm filter. Cells were fixed with 4% formaldehyde in phosphate buffered saline (PBS) and 0.05% Tween-80 for 30 minutes. Cells were washed and resuspended in PBS prior to analysis on the Amnis^®^ ImageStream^®^ X Mark II Imaging Flow Cytometer. A minimum of 20,000 in-focus events were captured per sample and the data were analysed using IDEAS 6.2 software. Signals from bright field, GFP and TurboFP635 were collected from channels 1, 2 and 4, respectively. The spot count feature was used to identify a population of single cells (identified as 1 object) after which the GFP best in focus population was selected for viable cells. A histogram of the normalized frequency against the intensity of TurboFP635 was generated for different pH conditions and time points and overlapped to demonstrate the formation of a viable, but non-replicating population.

VBNR bacteria can be identified by the retention of high TurboFP635. A gate was set on the top 50^th^ percentile of the induced TurboFP635 (pH 6.5 0h) culture, termed “high red”. This gate was used to determine the frequency of “high red” VBNR bacteria in pH 6.5 and pH 4.5 cultures at 48h. The percentage of VBNR subpopulation was calculated by the following equation:

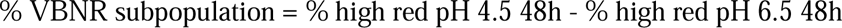

### Protein extraction

Following acid stress exposure of *M. tuberculosis* S169::pTiGc, the cell biomass was separated from the culture supernatant by centrifugation at 4000 rpm for 20 minutes.

Culture supernatants were sterilized by filtering twice using 0.22 µm Steriflip ® filter (Sigma Aldrich) and kept at under ice-cold conditions. Culture filtrates from pH 4.5 cultures were neutralized with sodium hydroxide to a pH of 6.5. All culture filtrates were concentrated using Amicon Ultra 3KDa spin columns (Sigma Aldrich) from 25 mL to 500 µL. SDS was added to a final concentration of 10% prior to incubation at 60°C for 1 hour and storage at -20°C.

Cell pellets from 25 mL cultures were stored at -20°C prior to resuspension in 1 mL lysis buffer (20 mM Tris-HCl, 1% SDS) containing protease inhibitors (Roche c0mplete^TM^ mini EDTA-free protease inhibitor cocktail). Cells were washed once for 2 minutes at 14 000 rpm, 4°C. Cell pellets were resuspended in 300 µL lysis buffer containing protease inhibitors and 10 µL DNAseI (RNAse-free) prior to cell disruption by mechanical bead-beating (Fast-prep-24^TM^, MPbio) using 300 µL acid-washed glass beads (425-600 µm, Sigma Aldrich). Bead-beating was performed 8 times for 20 seconds, at 4 m.s^-1^ with 1 minute intervals on ice. The cytosolic lysate was cleared by centrifugation at 12 000 rpm for 10 minutes at 4°C, the supernatant was recovered then clarified again by centrifugation as before. Clarified supernatants were filtered through 0.22 µm Acrodisks (Sigma Aldrich) and stored at -20°C.

### Mass spectrometry sample preparation

Culture filtrate and cell lysate proteins were reduced and alkylated prior to capture on MagReSyn^®^ HILIC beads (Resyn Biosciences). Briefly, the concentrated protein sample was reduced with 5 mM TCEP at room temperature for 1 hour and subsequently alkylated at 13 mM iodoacetamide for 1 hour in the dark. Iodoacetamide was quenched with 11 mM DTT. Cell lysate proteins were digested on MagReSyn^®^ HILIC beads as recommended by the manufacturer before storage at -20 °C.

Culture filtrate proteins were captured onto 150 µL prepared MagReSyn^®^ HILIC beads, overnight at 37°C. Beads were washed as per manufacturer recommendations and resuspended in 50 mM TEAB with 1 µg sequence modified trypsin. An on-bead digest was performed at 37°C for 18 hours, shaking at 800 rpm. Peptides were recovered and beads were incubated with 100 µL 1% TFA for 30 seconds at 800 rpm. The 1% TFA solution was recovered and pooled with recovered peptides from overnight digest. To ensure that bead captured proteins were digested, the protein digest was repeated with 100 µL TEAB and 1 µg sequence modified trypsin as before and recovered peptides were pooled and stored at -20°C.

Recovered supernatants were dried using a Concentrator*plus* (Eppendorf) and desalted as previously described (31). Peptide concentrations were determined against a 1 mg/mL peptide solution (Pierce) using a spectrophotometer (Jenway 7415).

### LC-MS/MS

Liquid chromatography was performed on an Ultimate 3000 RSLC equipped with a 20mm × 100μm C18 trap column (Thermo Scientific) and a CSH 25cm × 75μm, 1.7μm particle size C18 column (Waters). Solvent A (2% acetonitrile:water and 0.1% formic acid) was used to load samples onto the trap column from an autosampler (set to 7°C) at a flow rate of 2µL/min, for 5 minutes before sample elution onto the analytical column. A defined flow rate of 300 nL/min with the following gradient with Solvent B (100% acetonitrile, 0.1% formic acid) was used: 5–30% B over 60 min and 30–50% B from 60–80 min at 45°C.

Culture Filtrate samples were analysed in Data Dependent Acquisition (DDA) mode. Cell lysate samples were analysed in Data Independent Acquisition (DIA) mode.

DDA mass spectrometry analysis was performed using a Orbitrap Fusion^TM^ Tribird^TM^ Mass spectrometer (Thermo Scientific) equipped with a Nanospray Flex ionization source on positive mode with spray voltage set to 2 kV and ion transfer capillary at 290°C. Spectra were internally calibrated using polysiloxane ions at m/z = 445.12003. For MS1 scans the Orbitrap detector was set to a resolution of 60,000 over a scan range of 375–1500 with the AGC target at 4E5, and maximum injection time of 50 ms. Data was acquired in profile more. MS2 acquisitions were performed using monoisotopic precursor selection with ion charges +2 - +7 with an error tolerance of +/-10 ppm.

Precursor ions were excluded from repeat fragmentation for 60 s. Precursor ions were selected for fragmentation in HCD mode using the quadrupole mass analyser at an HCD energy of 30%. Fragments ions were detected in the Orbitrap mass analyzer with a resolution of 30,000. The AGC target was set to 5E4 and a maximum injection time of 60 ms. Data was acquired in centroid mode.

DIA mass spectrometry analysis was performed on the same instrument as for DDA analysis. The resolution of MS1 scans were set to 60,000 over a scan range of 375–1500. The AGC target was set to standard and a maximum injection time of 100 ms. Data was acquired in profile more. Precursor ions were selected for fragmentation in HCD mode using the quadrupole mass analyser at an HCD energy of 30%. Precursor ions were scanned in three windows, 355-555, 555-755, and 755-955 m/z and a 10 m/z isolation window with a 1 m/z overlap. Ions were detected in the Orbitrap mass analyzer set to 30,000 resolution and the AGC and maximum injection time set to custom. Data were acquired in centroid mode.

### Protein identification

MaxQuant 2.2.0.0 was used to analyze DDA tandem mass spectrometry data using the *M. tuberculosis* database (UP000001584) downloaded from Uniprot (https://www.uniprot.org/) in February 2023 (32,33). Carbamidomethyl cysteine was set as a fixed modification and oxidated methionine and N-terminal acetylation of proteins were selected as variable modifications. A maximum of 2 missed tryptic cleavages were allowed and proteins were identified with a minimum of 1 unique peptide. The protein and peptide false discovery rate (FDR) threshold was less than 0.01. Relative quantification was performed for identified protein groups using the MaxQuant LFQ algorithm and the “match between runs” algorithm was used to detect peptides which were not selected for MS/MS analysis in other replicate experiments.

Secreted proteins were identified using LFQ intensity data using Perseus (Figure 3). Potential contaminants, reverse hits, only identified by site potential contaminants, and proteins only identified with one unique peptide were removed. Proteins were considered true identifications if identified in at least two of the three biological replicate experiments for a particular condition. Proteins unique to either pH 6.5 or pH 4.5, where identifications were only made in one of the two conditions, were identified. For the remaining proteins, the LFQ intensity data was establish if proteins were significantly differentially abundant in the culture filtrates of pH 4.5 versus pH 6.5 cultures. The data was imputed by replacing missing values from the normal distribution of log2 transformed data for each experiment. Statistically significant differentially abundant proteins were identified following a paired student’s t-test and a Benjamini-Hochberg FDR correction of 0.05.

Cell lysate DIA data was analysed using FragPipe v22.0 (34–43). The DIA_SpecLib_Quant workflow was selected which built a spectral library using MSFragger-DIA against the *M. tuberculosis* UP000001584 database with added decoy and contaminant proteins using default settings. The library was filtered to 1% FDR at protein and peptide level. The spectral library was applied to quantify the DIA data using DIA-NN at 1% FDR. FragPipe-Analyst was used for the analysis and visualization of DIA data using the DIA workflow (44). Variance stabilizing normalization for DIA data was used for normalization and a Perseus-type imputation was selected for differential expression analysis. Differentially expressed genes were identified using a log fold change of 2 and an Benjamini Hochberg corrected p-value of 0.05. Protein groups only identified in a single biological replicate and contaminant protein groups were excluded from the identified protein list.

*M. tuberculosis* gene annotations, protein names and gene ontology information was obtained from Uniprot (http://www.uniprot.org/) and KEGG (https://www.genome.jp/kegg/) (33,45). Gene ontology enrichment analysis was done using the ShinyGO gene-set enrichment tool (https://bioinformatics.sdstate.edu/go/) (46).

## Results

### Acid stress promotes the formation of a viable, but non-replicating *M. tuberculosis* population

Clinical isolate *M. tuberculosis* S169 was obtained from a patient who remained culture positive following 6 months of TB treatment (27). Whole genome sequencing did not identify any known anti-TB treatment resistance conferring mutations and drug susceptibility was confirmed by phenotypic testing. Fluorescence dilution demonstrated an increased ability of this isolate to form VBNR *M. tuberculosis* in a macrophage infection model (7,28). We set out to establish if we could replicate the VBNR formation observed for *M. tuberculosis* S169 in a macrophage infection model using an *in vitro* low pH stress model (29). Briefly, *M. tuberculosis* S169 transformed with the replication reporter plasmid, pTiGc, was cultured in the presence of theophylline to induce the expression of TurboFP635. Cells were transferred into theophylline free culture media at either pH 6.5 or pH 4.5 and incubated for 48h (Figure 1A-B). Imaging flow cytometry demonstrated active replication of *M. tuberculosis* S169 at pH 6.5 with continued high levels of GFP expression, but reduced levels of red fluorescence following the removal of the inducer theophylline (Figure 1C). Following 48h of acid stress at pH 4.5, *M. tuberculosis* S169 continued to express high levels of GFP, but had reduced red fluorescence intensity, suggesting a decrease in replication (Figure 1C). Low pH stress induced the formation of 17.5% (+/- 4.5) VBNR *M. tuberculosis* S169, consistent with previous results investigating VBNR subpopulations using a macrophage infection model (7).

**Figure 1.**
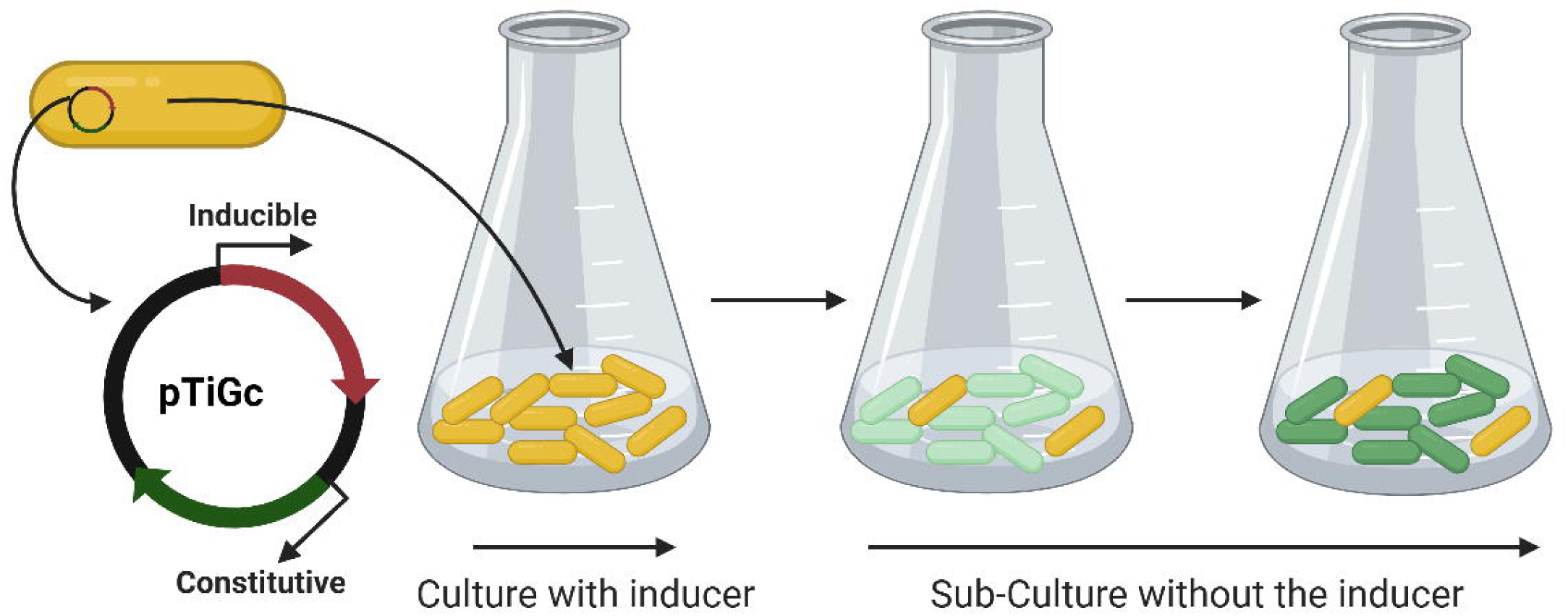
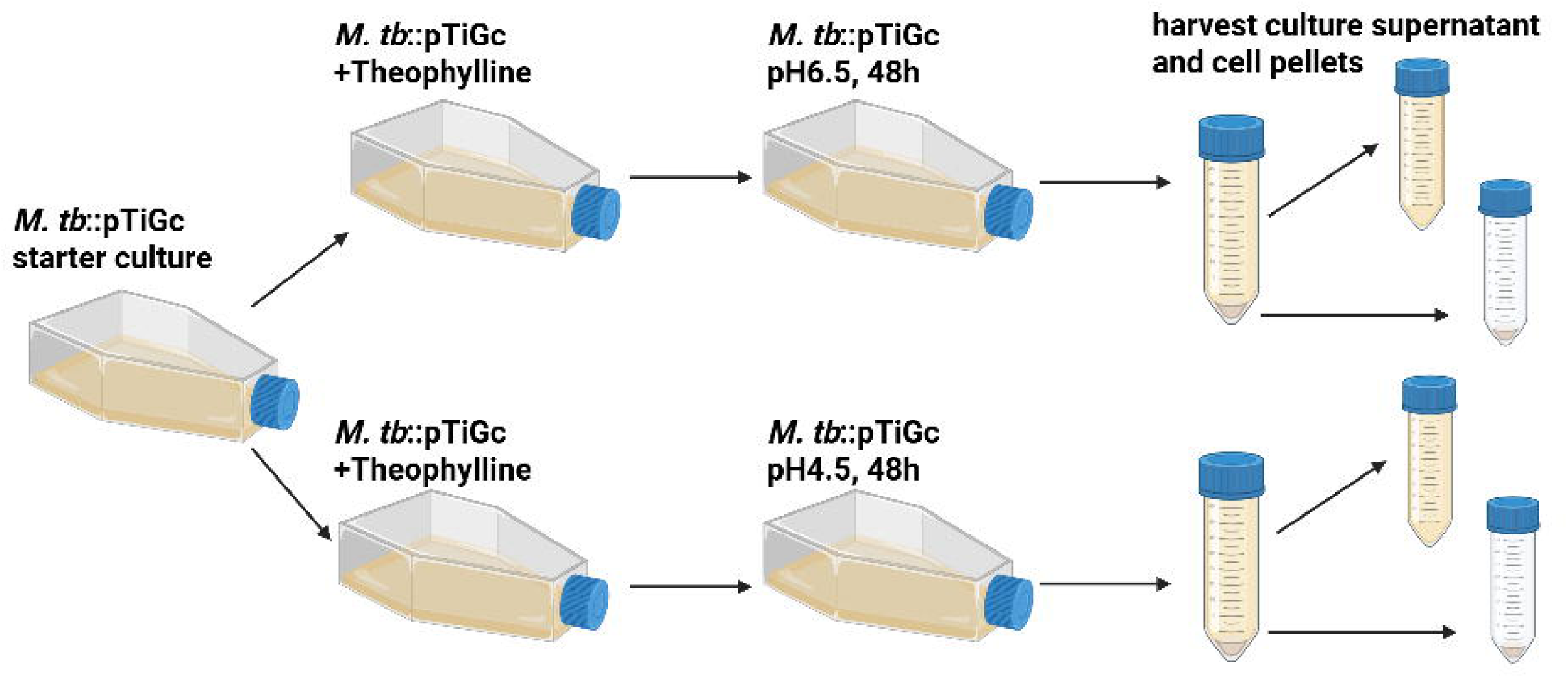
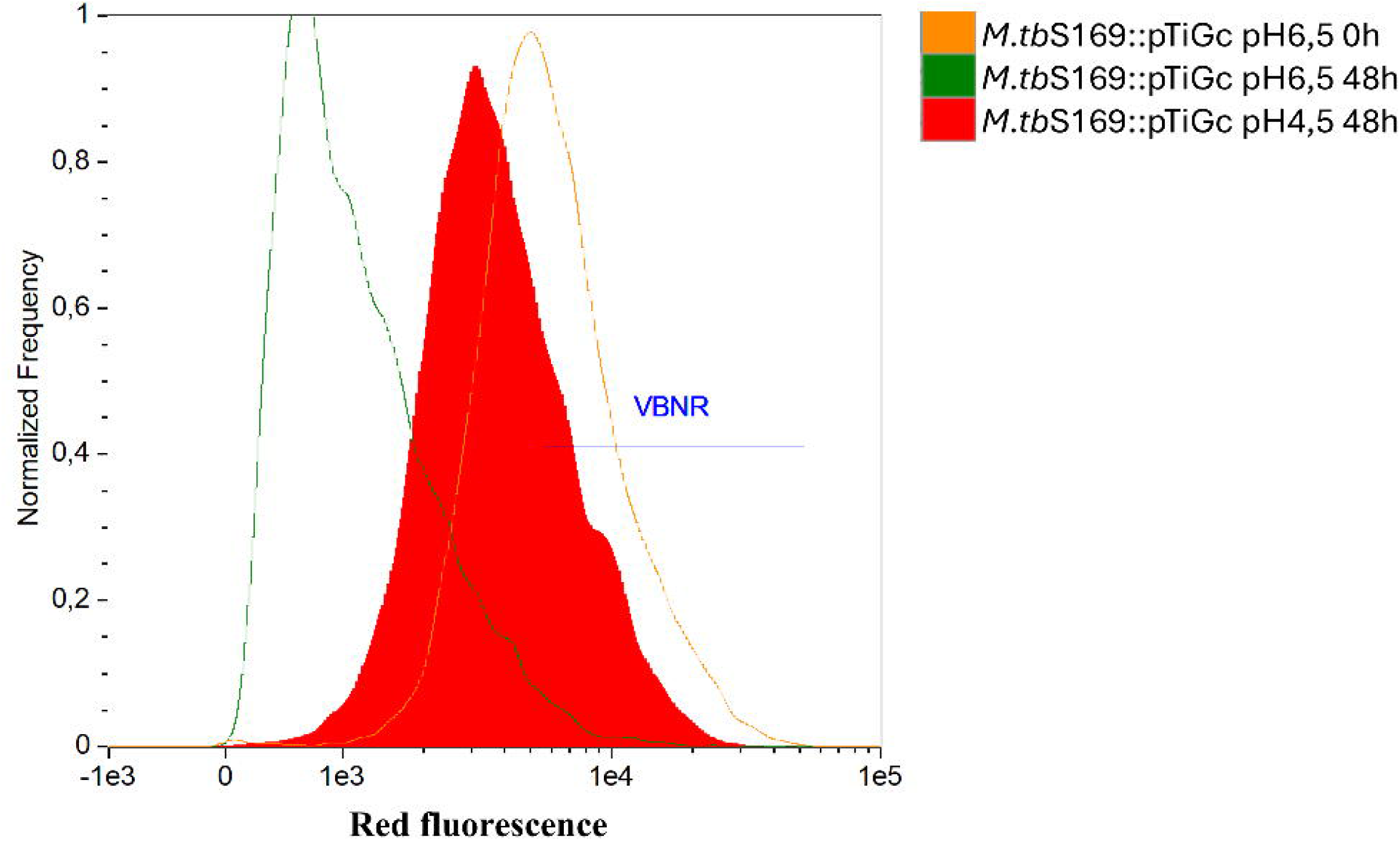
Acid stress promotes the formation of viable but non-replicating *M. tuberculosis*. **A.** *M. tuberculosis*::pTiGc will constitutively express a GFP (cell viability marker) and express TurboFP635 when cultured with theophylline (inducer). Upon removal of theophylline, VBNR or slowly replicating *M. tuberculosis* can be identified through retention of the red fluorescent signal. **B.** *M. tuberculosis* S169::pTiGc was cultured to an OD_600_ ∼ 1 prior to transferring into either pH6,5 or pH4,5 media for 48 hours. Aliquots of cultures were collected before and after stress for imaging flow cytometry. Cell pellets were collected for cell lysate proteomics and culture supernatants were collected for culture filtrate proteomics. **C.** *M. tuberculosis* S169::pTiGc was cultured with theophylline and a high intensity of red fluorescence was detected (orange). Following the removal of the inducer, actively replicating (pH6,5 cultures) bacteria had a reduction in red fluorescence (green), however, pH stressed cultures retained a high red fluorescence intensity (red). Created with BioRender.com.

### Low pH stress of *M. tuberculosis* S169 results in down-regulation of DosR

The cell biomass recovered from three independent low pH stress experiments was analysed using mass spectrometry to investigate the cellular stress response of the high VBNR *M. tuberculosis* S169 clinical isolate. We identified 2959 protein groups in the cell lysates of actively replicating and acid stressed *M. tuberculosis* S169 which mapped to 2924 proteins in the KEGG database (Table S1) following the removal of contaminant proteins and proteins only identified in a single biological replicate. A comparison of actively replicating and low pH stressed cell lysates revealed that 46 proteins were only identified in the cell lysates of actively replicating *M. tuberculosis* S169 (Table S2) and an additional 14 proteins were only identified in the cell lysates of VBNR-enriched *M. tuberculosis* S169 (Table S3). Differential analysis of the cell lysate data revealed that 77 proteins were significantly more abundant, and 269 proteins were significantly less abundant (adjusted p-value <0,05, log2 fold change >1) in the cell lysates of low pH stressed *M. tuberculosis* S169 when compared to that of actively replicating *M. tuberculosis* S169 (Table S1, Figure 2A).

**Figure 2.**
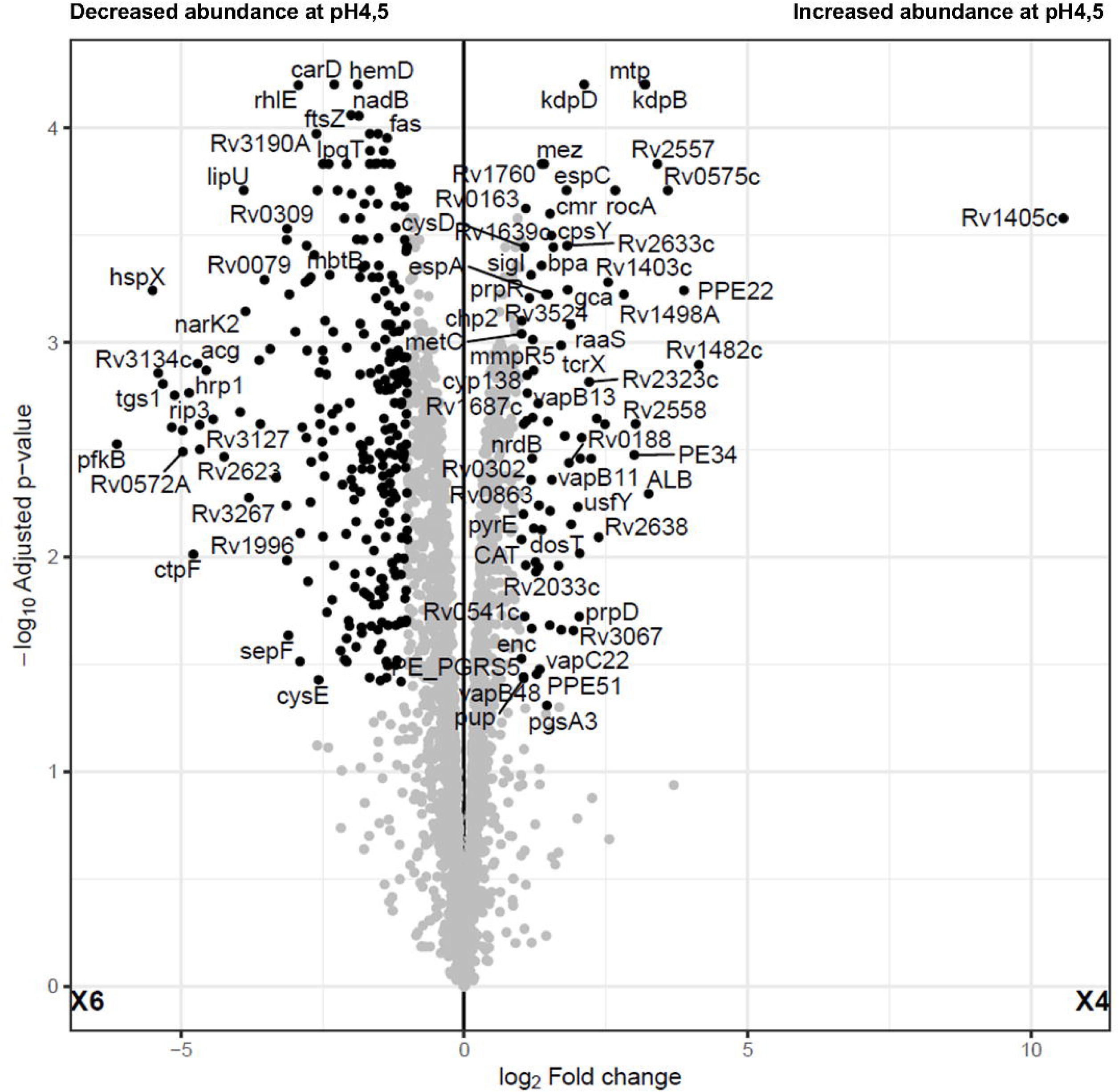
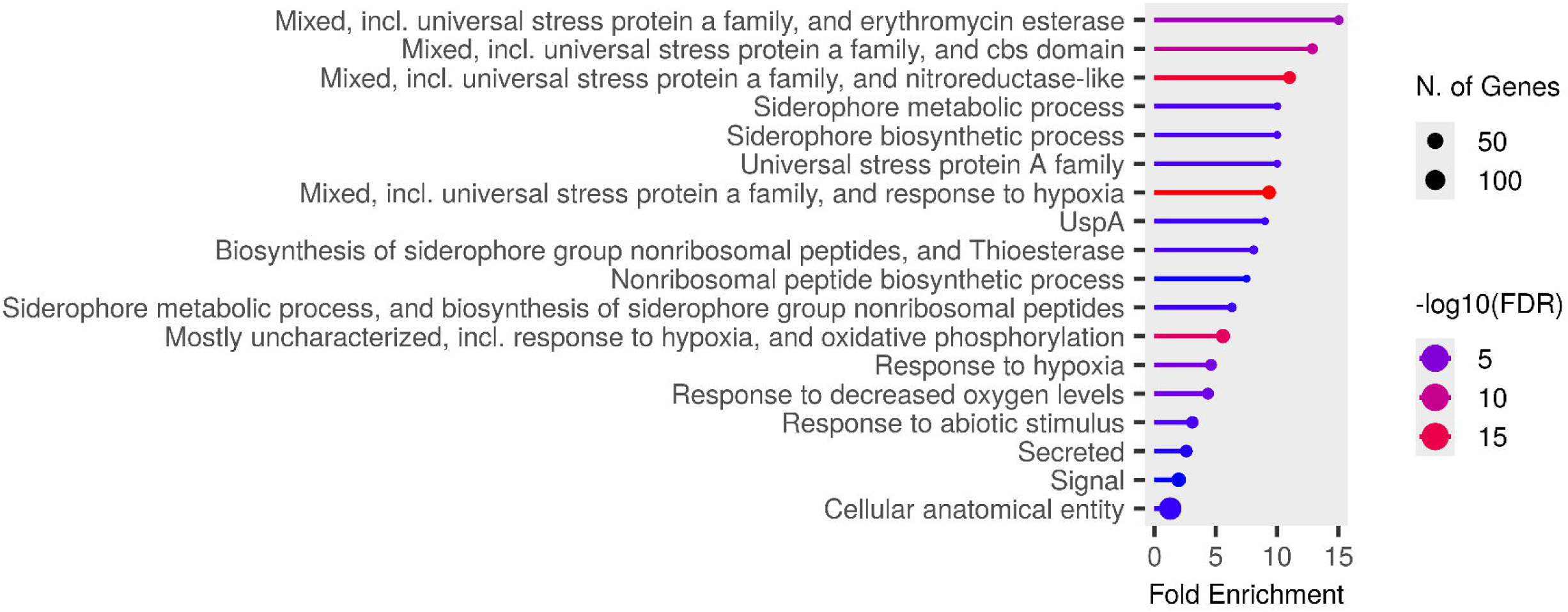
Proteins identified in the cell lysates of actively replicating and VBNR enriched cell lysates. **A.** Volcano plot representing differential protein abundance for *M. tuberculosis* S169 cell lysate proteins. Volcano plot generated by FragPipe-Analyst using protein abundance data for proteins identified in the cell lysates of actively replicating and acid stressed *M. tuberculosis* S169. Following the removal of contaminant proteins and proteins only identified in one biological replicate, 77 proteins were more abundant, and 269 proteins were less abundant in pH 4.5 cell lysates when compared to pH 6.5 cell lysates (adjusted p-value <0,05, fold change >2). **B.** Gene ontology enrichment of proteins significantly less abundant in the cell lysates of acid stressed *M. tuberculosis* S169. GO enrichment analysis revealed that GO terms associated with universal stress proteins were enriched in the 269 proteins found to be significantly less abundant in the cell lysates of acid stressed *M. tuberculosis* S169 when compared to that of actively replicating *M. tuberculosis* S169.

Gene ontology (GO) enrichment analysis of proteins significantly more abundant in the cell lysates of acid stressed cultures did not reveal any significant results. Regardless, the two-component system TcrXY component TcrX was significantly more abundant in acid stressed *M. tuberculosis* S169 (Table S1). The TcrXY two component system has previously been shown to be upregulated in response to acid stress (47). S-adenosyl methionine (SAM)-dependent methyltransferase proteins Rv1403c and Rv1405c were also significantly more abundant in acid stressed *M. tuberculosis* S169 (Table S1). These SAM-dependent methyltransferases have previously been reported to be upregulated in response to low pH (48–50).

A GO enrichment analysis of significantly less abundant proteins in cell lysates of acid stressed cultures suggested a down regulation of GO terms associated with universal stress response proteins (Figure 2B, Table S4). The dormancy response regulon, under the control of the two-component system DevR/DevS, has previously been shown to be upregulated in response to acid stress (29,51). The DosR regulon is composed of 47 genes (52). In our study, we identified 38 DosR regulon encoded proteins of which 34 were significantly less abundant in the cell lysates of acid stressed *M. tuberculosis* S169 (Table 1).

**Table 1.**
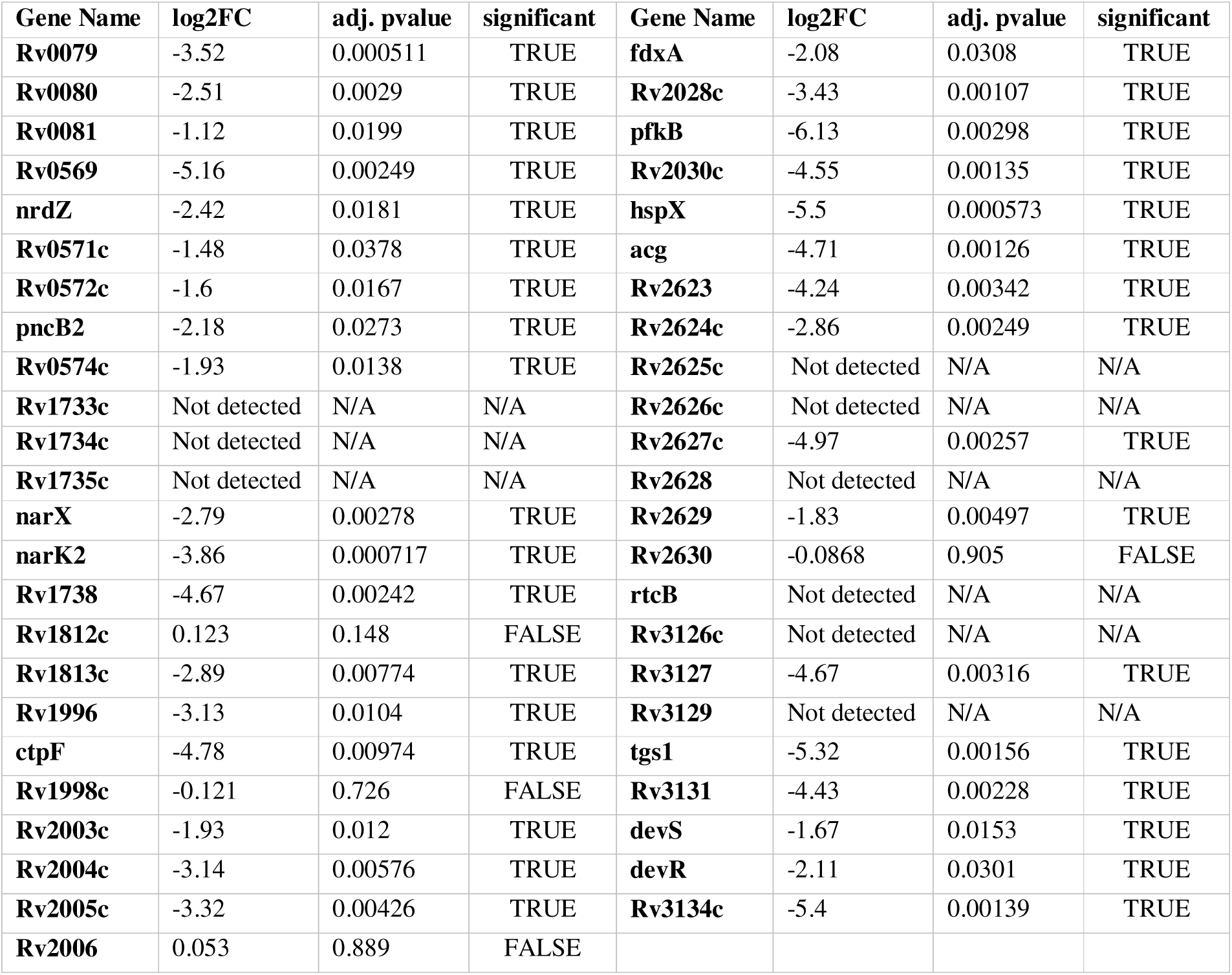
Abundance of DosR proteins in *M. tuberculosis* S169 in response to acid stress.

### Culture filtrates of VBNR enriched cultures showed a higher abundance of lipoproteins

In total, we identified 461 protein groups of which 192 protein groups were only identified in the culture filtrates of pH 6.5 cultures and 45 protein groups were only identified in the culture filtrates of pH 4.5 cultures (Figure 3). Abundance data of protein groups identified in both test conditions revealed that 83 protein groups were differentially abundant between the conditions tested (q-value < 0.05, FDR 0.05, log2 FC >/<0.05) (Figure 3, Table S5-6). Of the 83 significantly differentially abundant proteins, 43 protein groups were less abundant, and 40 proteins were more abundant in the culture filtrates of VBNR enriched *M. tuberculosis* S169 (Table S5-6). In total, we identified 275 protein groups, which mapped to 274 proteins in the KEGG database, in the culture filtrates of actively replicating *M. tuberculosis* S169 cultures (Table S5). The 128 protein groups identified in the culture filtrates of VBNR-enriched cultures mapped to 128 proteins in the KEGG data base (Table S6).

**Figure 3.**
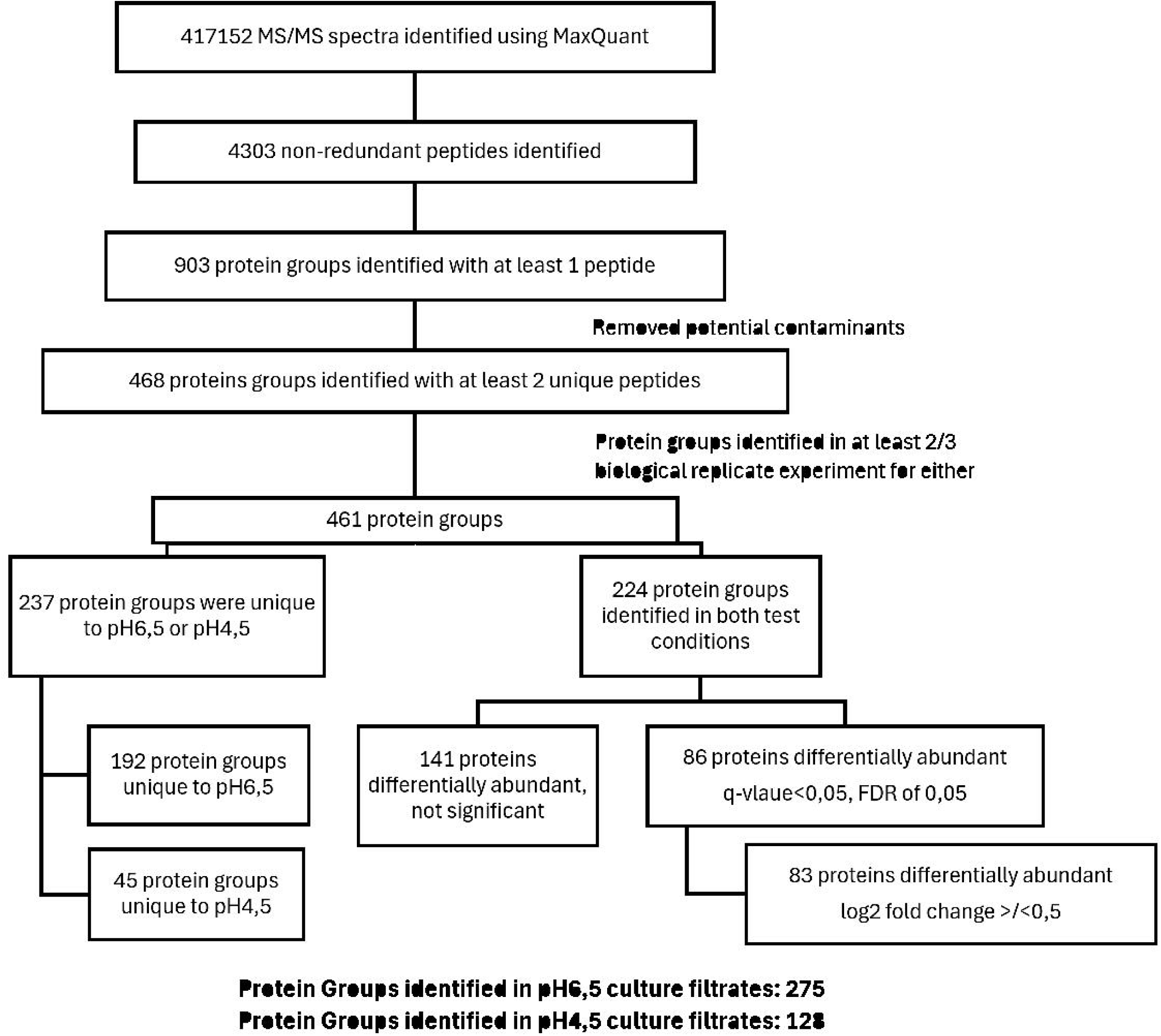
Identification of culture filtrate proteins from pH 6.5 and pH 4.5 cultures. The diagram demonstrates the data analysis of DDA mass spectrometry data for *M.* tuberculosis S169 culture filtrates. Following automated database searching, 461 protein groups were identified in at least two of the three biological replicate experiments in culture filtrate recovered from pH 6.5 and pH 4.5 cultures. Of these, 192 protein groups were only identified in pH 6.5 cultures filtrates and 45 proteins were only identified in pH 4.5 culture filtrates. For proteins identified under both test conditions, a paired student t-test with a Benjamini-Hochberg correction of 0,05 was used to identify 83 differentially abundant proteins (log2 FC >/<0,05). In total we identified 275 protein groups in the culture filtrates of pH 6.5 culture and 128 protein groups in the culture filtrates of pH 4.5 cultures.

GO enrichment of proteins identified in the culture filtrates of actively replicating and VBNR enriched *M. tuberculosis* S169 cultures confirmed the enrichment of pathways extracellular region, external encapsulating structure, and secreted (Figure 4, Table S7-8). Other enriched pathways included cell wall, cell periphery, plasma membrane, and membrane (Figure 4, Table S7-8). The lipoprotein pathway was revealed to be enriched in the culture filtrates of pH 4.5 cultures (Figure 4B, Table S8). Several lipoproteins were only identified in VBNR enriched culture filtrates (LpqG, DppA, LpqO, FecB2, LppL, LppM, Subl, GlnH, and Rv2585c) or identified with a higher relative abundance in pH 4.5 culture filtrates (FecB, LpqB, LprG, and LprA) (Table S6). Zymogen binding, preceding the proteolytic cleavage of enzymes to an active state, was also enriched in the culture filtrates of acid-stressed *M. tuberculosis* S169 (Figure 4B). Zymogen binding proteins were more abundant in the culture filtrates of VBNR enriched cultures, including MetK, LpdC, Mpt64, GroES, FbpA and FpbB (Table S6, S8). Proteases were also enriched within the culture filtrates of acid stressed *M. tuberculosis* S169 and included proteases HtrA1, PepA, PepD, Rv3671c, Clp1, Clp2, MycP3 and Rv2672 (Table S6, S9).

**Figure 4.**
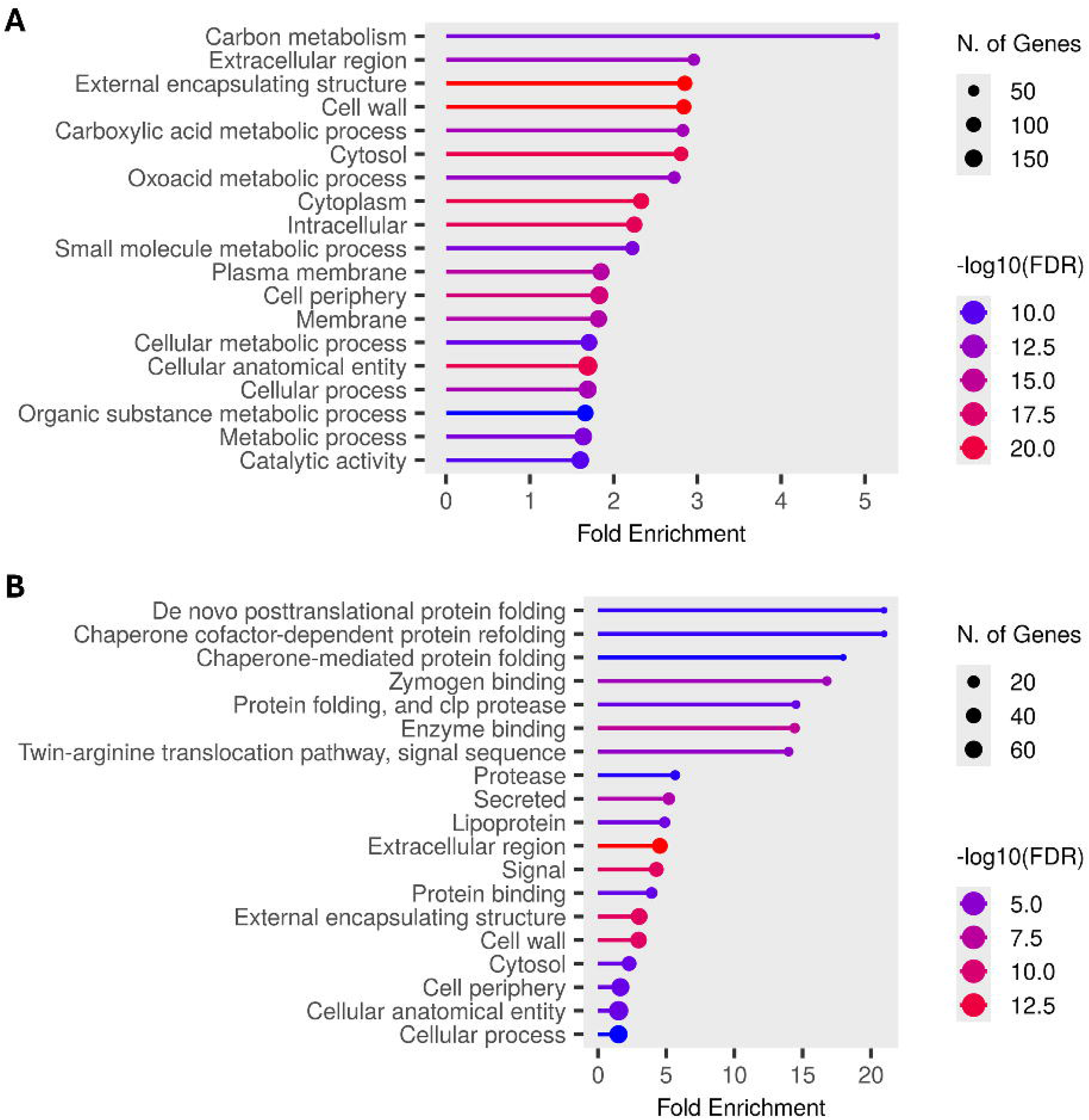
Enriched pathways of proteins identified in the culture filtrates of actively replicating and acid stressed *M. tuberculosis* S169. GO enrichment analysis confirmed the enrichment of proteins from the extracellular region through enrichment of pathways secreted and extracellular region. **A.** GO enrichment analysis of proteins from actively replicating *M. tuberculosis* S169 revealed the enrichment of pathways for carbon metabolism, carboxylic acid metabolism, and oxoacid metabolic process. **B.** Proteins identified in the extracellular fraction of acid-stressed *M. tuberculosis* S169 revealed the enrichment of pathways for protein folding, lipoprotein, zymogen and enzyme binding, and protease.

## Discussion

Phagosome acidification is an environmental stress faced by *M. tuberculosis* during host infection (53). In this study, we took advantage of a low pH stress model to trigger the formation of a *M. tuberculosis* VBNR subpopulation (29). To increase the probability of identifying VBNR secreted proteins, we made use of a clinical isolate, *M. tuberculosis* S169, which we previously showed to form high proportions of VBNR bacteria (7,27). The fluorescence dilution replication plasmid enabled the tracking of bacterial replication in response to low pH stress (Figure 1A) (28,29). Following acid-stress for 48h at pH 4.5 (Figure 1B), imaging flow cytometry confirmed the formation of a large VBNR subpopulation of *M. tuberculosis* S169 (Figure 1C).

Proteomic characterization of the cell lysate revealed the differential abundance of 346 proteins in response to acid-stress (Figure 2A, Table S1). Of the 77 proteins significantly more abundant (adjusted p-value <0,05, fold change >2) in the cell lysates of acid stressed *M. tuberculosis* S169, no enriched pathways were identified using GO enrichment analysis. However, in agreement with previous reports, the TcrXY two-component system components were more abundant in acid-stressed *M. tuberculosis* S169, with the response regulator TcrX significantly more abundant (Table S1) (47). TcrX is required for *M. tuberculosis* survival during chronic infection (47). Similarly, the acid stress-induced methyltransferases Rv1403c and Rv1405c were also significantly more abundant in acid stressed *M. tuberculosis* S169 (Table S1), as previously reported for *M. tuberculosis* (49,50,54). Interestingly, Rv1405c has also been reported to be upregulated during the enduring hypoxic response and nitrosative stress (55,56). Even though Rv1405c is not essential for *in vitro* survival, it is required for survival in C57BL/6J mice (57,58). The role of the Rv1405c methyltransferase during infection remains unknown.

Two-component systems are required by the bacteria to respond to environmental changes. The PhoP component from the PhoPR two-component system is known to positively regulate the *aprABC* operon in response to acidic pH (59–61). In this study, the AprA protein was only identified in acid stressed *M. tuberculosis* S169 (Table S1, S3) and AprB and AprC proteins were not detected (Table S1). Interestingly, despite detection of AprA in VBNR enriched cultures, PhoPR components were found to be less abundant in the cell lysates of acid stressed bacteria (Table S1). The two-component system, KdpD/KdpE, has been suggested to play a role in the evasion of phagocytic killing and enabling bacterial persistence (62). In this study, KdpA, KdpB and KdpD were all significantly more abundant in the cell lysates of VBNR-enriched cultures (Table S1). KdpE was found to be more abundant, but not significantly (Table S1). Interestingly, KdpA has been suggested to be required for ATP homeostasis and persister formation in *Mycobacterium marinum* (63).

In response to acid stress, 269 cell lysate proteins were significantly less abundant (adjusted p-value <0,05, fold change >2) than in the of actively replicating *M. tuberculosis* S169 (Table S1). Gene Ontology enrichments revealed a lower abundance of universal stress proteins (Figure 2B, Table S4), including components from the two-component system DosR, also known as the dormancy response regulon. DosR is known to be upregulated in response to hypoxia, starvation and low pH (29,51,64). Interestingly, in this study, 34 components of the DosR regulon were significantly less abundant in the acid-stressed *M. tuberculosis* S169 (Table 1). These results contrast with some previous reports, which largely show an upregulation of DosR in response to environmental stress. However, a lower induction for DosR in response to low pH has been reported in comparison to hypoxia, starvation and stationary phase growth (51). In agreement with our results, DosR components Rv0080, NarX, Rv2030c, and Rv1813c have previously been shown to be downregulated in response to acid stress (51). The DosR regulon is largely down regulated in a *M. tuberculosis* pellicle biofilm model (65). More recently, another pellicle biofilm study showed the down regulation of DosR genes in five of the six *M. tuberculosis* lineage 4 clinical isolates studied (66). The clinical isolate investigated in this study, *M. tuberculosis* S169, belongs to lineage 4 (7). Interestingly, *M. tuberculosis* H37Rv, in which the DosR regulon is upregulated in response to low pH, also belongs to lineage 4 (29). These findings highlight the need to investigate the response of *M. tuberculosis* clinical isolates to physiologically relevant stress conditions for a more comprehensive understanding of the mycobacterial stress response.

RocA, EspA, EspC, Rv2390c, and PE34 were significantly more abundant in the cell lysates of acid stressed *M. tuberculosis* S169, as previously reported (Table S1) (67). ESX-1 is important for *M. tuberculosis* virulence and EspA and EspC are ESX-1 secretion associated proteins (68–71). Several other ESX-1 proteins were more abundant in the cell lysates of acid stressed cultures (Table S1) (72–74). Despite the increased abundance of EspA, EspB, EspD in acid stressed cell lysates, these proteins were not detected in the culture filtrates of acid stressed *M. tuberculosis* S169 (Table S6). EspF, EspC, EspH, EspR, and EspK proteins were present in the culture filtrates of actively replicating *M. tuberculosis* S169 (Table S5). In agreement with previous reports, Rv0516c, LipL, and PPE59 were less abundant in response to acid stress (67). Interestingly, PPE22 has not previously been reported to be upregulated in response to acid stress, however, in this study PPE22 was significantly more abundant in the cell lysates of acid stressed *M. tuberculosis* S169 (Table S1) (67). Moreso, PPE22 was only identified in the culture filtrates of acid stressed cultures (Table S6). PPE22 has been previously been detected in guinea pig lungs at 30 days post infection and more recently has been shown to induce a protective immune response in BALB/c mice, showing promise as a vaccine development candidate (75,76).

Several cell division proteins were less abundant in the cell lysates of VBNR-enriched cultures including WhiB2, MtrA, SepF, FtsZ, FtsK, FtsQ, FtsW, FtsE, CwsA, and CrgA (Table S1). We also detected a lower abundance of DNA replication and repair proteins, including ImuA, RecA, RecR, RecN, DnaB, and Rv1277 (Table S1). The downregulation of DNA replication and repair proteins and cell division protein aligns with reduced bacterial replication, as observed with the fluorescence dilution experiments (Figure 1). Low pH stress induced the formation of a VBNR subpopulation (Figure 1C). Interestingly, resuscitation-promoting factors (Rpf) RipA, RipB, and RpfC were significantly less abundant in the cell lysates of VBNR enriched *M. tuberculosis* S169 cultures (Table S1). RipB was detected in the culture filtrates of both actively replicating and VBNR enriched cultures, but at a significantly lower abundance in VBNR enriched culture filtrates (Table S5-6). Other Rpf proteins were identified in the culture filtrates of both test conditions, but with no significant difference in abundance (Table S9). Rpf muralytic enzymes stimulate growth of dormant *M. tuberculosis* and the loss of Rpfs results in an impaired ability of *M. tuberculosis* to resuscitate from a non-culturable state (77).

Culture supernatants have low protein concentrations, often resulting in the need to pool culture supernatants from multiple cultures to a obtain enough protein. This practice increases the possibility of introducing inter-culture variation. To overcome this limitation, we applied a protein aggregation capture approach to study the culture supernatant of a single bacterial culture per replicate experiment. A single culture has the benefit of reducing time, cost, and biohazardous risk in addition to limiting technical and biological variation from multiple cultures. Applying this approach, we showed that the culture filtrates of actively replicating *M. tuberculosis* S169 contained 274 proteins compared to the 128 proteins identified in the culture filtrates of VBNR enriched *M. tuberculosis*. GO pathway enrichment analysis confirmed the enrichment of extracellular region and secreted pathways (Figure 4). Zymogen binding, lipoprotein, protein folding, and protease pathways were enriched from proteins identified in the extracellular fraction of low pH stressed *M. tuberculosis* S169 (Figure 4B). Zymogens are the inactive precursors of enzymes which get converted to active forms by proteolysis. Lipoproteins have been implicated in *M. tuberculosis* virulence and immune modulation (78). Other enriched pathways included the external encapsulating structure, cell periphery, and plasma membrane which may be the result of culturing *M. tuberculosis* S169 in media containing the detergent Tween-80 to prevent bacterial clumping. The inclusion of Tween-80 in *M. tuberculosis* culture media has been speculated to result in the solubilization of lipids and the shedding of surface adhered molecules (79,80). As indicated by the pathway enrichment analysis, cytosolic proteins including RNA polymerase subunits were identified in actively replicating *M. tuberculosis* S169 culture filtrates (Table S5). Small ribosomal subunits were also identified in the culture filtrates of both actively replicating and VBNR enriched *M. tuberculosis* S169 cultures (Table S5 and S6). The identification of these proteins outside the cell may suggest some cell lysis occurred. The culture filtrates of VBNR enriched *M. tuberculosis* S169 cultures contained 45 proteins not identified in the extracellular fraction of actively replicating bacteria (Table S6). These proteins and the 83 significantly differentially abundant proteins (40 more abundant and 43 less abundant) identified in the culture filtrates of acid stressed *M. tuberculosis* S169 suggest that *M. tuberculosis* may secrete a different subset of proteins in response to low pH stress. The VBNR subpopulation only accounted for 17.5% (+/- 4.5) of the bacterial population investigated at pH 4.5, however, we speculate that VBNR protein secretion contributed to the differences observed in the culture filtrates between pH 6.5 and pH 4.5 cultures. Although not investigated in this study, differences in the proteins found in the extracellular region of *M. tuberculosis* S169, may result in a different immune response during infection. Culture filtrates from VBNR enriched cultures included proteins from Toxin-antitoxin (TA) systems, VapC51 and VapB10 (Table S6). TA systems have been implicated in the adaptation to environmental stress and bacterial persistence. Type II TA systems are highly abundant in *M. tuberculosis* genomes (10,81). Interestingly, the chorismate mutase Rv1885c was only identified in the secreted fraction of VBNR enriched cultures, and was recently suggested contribute to *Mycobacterium bovis* BCG pathogenesis by inhibiting mitochondria-mediated cell death of macrophages (82). Immunogenic proteins more abundant in the culture filtrates of VBNR enriched cultures included FbpA (Mpt44), Mpt53, Mpt64 and Mpt63 (Table S6).

In this study we investigated the cellular proteome and the extracellular region of a clinical isolate with an increased propensity to form VBNR bacteria in response to low pH stress (7). We acknowledge that our study was limited by only investigating a single clinical isolate, however, this isolate was chosen to increase the likelihood of identifying changes in the proteome because of its increased propensity to form VBNR bacteria. We demonstrated that this clinical isolate did form a viable but non-replicating population in response to *in vitro* low pH stress. Cell lysate proteomics revealed increased abundance of known acid stress proteins, however, in contrast to what has been published previously, several proteins of the DosR response regulon were significantly less abundant in low pH stressed *M. tuberculosis* S169. This study highlights the need to investigate the cellular response of clinical isolates, specifically clinical isolates obtained from individuals with unfavourable outcomes, to improve our understanding of factors which may contribute to treatment failure. Using our culture filtrate mass spectrometry approach, we demonstrated that the culture filtrate composition of actively replicating and low pH stressed VBNR enriched cultures had different compositions. While we cannot definitively demonstrate secretion of proteins by VBNR bacteria, several proteins identified in the culture filtrates of VBNR enriched cultures have implicated roles in bacterial persistence. Importantly, the culture filtrate approached used in this study has the potential to be used not only to investigate *M. tuberculosis* extracellular fractions but can be adapted to study extracellular proteins in other bacteria.

## Declarations

### Ethics approval statement

Ethics approval was obtained from the Human Research Ethics Committee (N10/01/013) and the Biological and Environmental Safety Committee (BES-2023-13049) at Stellenbosch University.

### Consent for publication

Not applicable.

### Availability of data and materials

Imaging flow cytometry data are available from the corresponding author upon request. Mass spectrometry proteomics data are available from the ProteomeXchange Consortium via the PRIDE partner repository with the identifiers PXD068623 and PXD068720 (83).

### Funding

This research was supported by the VALIDATE Network which was funded by Gates Foundation (INV-031830) and the South African government through the National Research Foundation of South Africa (NRF) and the South African Medical Research Council (SAMRC). NK acknowledges research and salary support from the VALIDATE Network, which was funded by the Gates Foundation (INV-031830). SS is funded by the South African Research Chairs Initiative of the Department of Science and Technology and National Research Foundation (NRF) of South Africa, award number UID 86539.

The authors are all affiliated with the with the DSI-NRF Centre of Excellence for Biomedical Tuberculosis Research; South African Medical Research Council Centre for Tuberculosis Research; Division of Molecular Biology and Human Genetics, Faculty of Medicine and Health Sciences, Stellenbosch University, Cape Town.

### Authors contributions

NK, JC, JM, and SS assisted with experimental design and conceptualization. NK and JC performed the experimental work and NK analyzed the results. NK drafted the manuscript, tables, and figures. All authors contributed to this manuscript and approved the submitted version.

## Acknowledgements

We acknowledge Maré Volk from the Central Analytical Facilities at Stellenbosch University for technical assistance for mass spectrometry.

Figure 1A was created in BioRender. Sampson, S. (2025) https://BioRender.com/149jzej.

Figure 1B was created in BioRender. Sampson, S. (2025) https://BioRender.com/fyvj3w9.

